# The effect of crack cocaine addiction and age on the microstructure and morphology of the human striatum and thalamus using shape analysis and fast diffusion kurtosis imaging

**DOI:** 10.1101/066647

**Authors:** Eduardo A Garza-Villarreal, M Mallar Chakravarty, Brian Hansen, Simon F Eskildsen, Gabriel A. Devenyi, Diana Castillo-Padilla, Thania Balducci, Ernesto Reyes-Zamorano, Sune N Jespersen, Pamela Perez-Palacios, Raihaan Patel, Jorge J Gonzalez-Olvera

## Abstract

The striatum and thalamus are subcortical structures intimately involved in addiction. The morphology and microstructure of these have been studied in murine models of cocaine addiction, showing an effect of drug use, but also chronological age in morphology. Human studies using non-invasive MRI have shown inconsistencies in volume changes, and have also shown an age effect. In this exploratory study, we used MRI-based volumetric and novel shape analysis, as well as a novel fast diffusion kurtosis imaging sequence to study the morphology and microstructure of striatum and thalamus in crack cocaine addiction (CA) compared to matched healthy controls (HC), while investigating the effect of age and years of cocaine consumption. We did not find significant differences in volume and mean kurtosis (MKT) between groups. However, we found significant contraction of nucleus accumbens in CA compared to HC. We also found significant age related changes in volume and MKT of CA in striatum and thalamus that are different to those seen in normal aging. Interestingly, we found different effects and contributions of age and years of consumption in volume, displacement and MKT changes, suggesting each measure provides different but complementing information about morphological brain changes and that not all changes are related to the toxicity or the addiction to the drug. Our findings suggest that the use of finer methods and sequences provide complementing information about morphological and microstructural changes in cocaine addiction, and that brain alterations in cocaine addiction are related cocaine use and age differently.

## Introduction

The striatum and thalamus are subcortical structures greatly affected in the physiopathology of cocaine addiction in animal models and possibly in humans ^1^. Studies in animal models show microstructural changes in striatal medium spiny neurons, as well as striatal indirect, direct, and thalamo-striatal pathways ^2^. Using neuroimaging in mice, Wheeler *et al* ^3^ showed volumetric reduction in ventral and posterior striatum, and expansion in the dorsal striatum in a cocaine addiction model. However, non-invasive neuroimaging studies in human cocaine addicts have shown inconsistent and conflicting neuroanatomical abnormalities in striatum and thalamus ^4,5^.

A number of studies have observed lower volume in striatum ^6,7^, larger striatal and thalamic volumes ^8–10^, reduced volume in anterior and increase in posterior striatum ^11^, or no volumetric differences whatsoever^12,13^. These discrepancies may be explained by methodological factors (i.e. the use of volume as a metric, the segmentation method, small sample sizes), but also, polysubstance use and cocaine delivery (inhaled vs smoked) could play an important a role. Compared to inhaled cocaine, crack cocaine (smoked) is clinically related to stronger craving, addiction and deterioration in cognition^14,15^. Crack cocaine use and addiction is also most prevalent in the lower socioeconomic stratum and poses a great health risk to addicts both directly and indirectly, as well as being a great social problem and expense ^16,17^.

To better understand the neuroanatomical effects of cocaine as the main drug of use and the mechanism of addiction, there is a need for large-scale studies using more sophisticated computational neuroanatomical methods and novel neuroimaging sequences in human addicts, while focusing on one type of cocaine delivery separately. Also, it would be important to study brain morphology in a wider range of ages (young and older addicts), as it has been shown that brain changes related to cocaine abuse tend to worsen with age in young mice ^3^ and in humans ^18^. Because substance addiction is known to affect both the brain gross anatomy (volume and shape) and tissue composition (microstructure), our study employs an imaging protocol allowing both types of analysis. Volumetric and shape analysis are performed based on high-resolution T1-weighted magnetic resonance imaging data. Shape analysis refers to the study of the three dimensional shape of the subcortical nuclei using the information of the automatic segmentation and shape-based models ^19^. The average nonlinear deformations are used to estimate displacement of the nuclei with respect to the model (inward or contraction, outward or expansion displacement), as well as surface area calculated from a tessellated surface of the segmentation. These metrics, surface area and displacement, are related to biological processes and pathology (e.g. schizophrenia), and provide different and complementary information than volume ^20,21^.

Diffusion kurtosis imaging is a novel imaging technique known to be very sensitive to tissue microstructure, even in regions of crossing fibers. However, it has not been used for the study of substance abuse and addiction. The basis of DKI is that in tissue, microstructural components hinder free (i.e. Gaussian) diffusion of water ^22^. The measured diffusion MRI signal yields this deviation from normal diffusion as a non-zero kurtosis ^23^, providing an indirect measure more sensitive to the microstructural physical composition than fractional anisotropy or other diffusion tensor based methods. DKI, however, may or may not be related to macrostructural volume or shape changes measured with T1w images, as DKI measures microstructural changes. One typically reported DKI metric is the mean kurtosis (MK) which has been associated with microstructural changes in a host of diseases such as stroke ^24^, but has also been shown to be sensitive to more subtle brain alterations as in mild traumatic brain injury ^25,26^, mild chronic stress ^27^, and even the brain remodeling that is part of normal aging ^28^. For this study we employed a fast DKI (fDKI) sequence allowing rapid acquisition of the data needed for estimation of the tensor-based mean kurtosis ^29–31^. The fDKI method uses a tensor-based mean kurtosis definition (MKT) that differs slightly from the traditional mean kurtosis (MK), nevertheless the agreement between MK and the rapidly obtainable MKT is well-established ^32^ and clinical applications are favorable ^33^. In our exploratory study we compare the striatum and thalamus morphology and microstructure of active crack cocaine addicts with healthy controls, using novel analytic computational neuroanatomical methods and a recently introduced fast DKI sequence ^30^. We also included a post-hoc striatum and thalamus subdivision analysis due to the importance of the different subnuclei in the pathology such as the ventral striatum ^1,34^.

## Materials and Methods

### Participants

We recruited 54 cocaine addicts and 48 healthy controls (HC) from March to December of 2015 as part of a principal addiction project. Healthy controls were matched by age (± 2y), sex and handedness. Education was matched as closely as possible. The recruitment criteria are shown in Supplementary Table 1. We decided to only study a subset of crack cocaine addicts (CA) for three reasons: 1) The known difference in terms of effects and dependency between inhaled and smoked cocaine ^15^, 2) the small sample size of our inhaled cocaine addicts before exclusion and elimination (n = 13), and 3) crack cocaine is a more important socio-economical issue in Mexico and the world due to the toxicity of it, the stronger craving and the low social stratum where it is mostly used. We then excluded 9 participants (HC = 6, CA = 3), eliminated 1 participant due to claustrophobia (HC) and 3 due to excessive movement during image acquisition (HC = 1, CA = 2). Our final sample size for the morphological analysis was 36 CA (3F) / 40 HC (2F) with a median age of 30 (18 – 48) years old. For the fDKI analysis, we had a final sample size of 17 CA / 18 HC as the DKI sequence was acquired in a smaller subset of participants and due to the outlier rejection using Thompson’s Tau ^35^.

The study was approved by the local ethics committee and performed at the Instituto Nacional de Psiquiatría “Ramón de la Fuente Muñiz” in Mexico City, Mexico. The study was carried out according to the Declaration of Helsinki. All participants were invited through posters placed in several centers for addiction treatment. Healthy controls were recruited from the Institute (i.e. administrative workers) and using fliers around the city. Participants had the study fully explained to them and provided verbal and written informed consent. The participants underwent clinical and cognitive tests besides the MRI as part of the main ongoing addiction project (see Supplementary Methods and Results and Supplementary Table 2 for details), which are not part of this study. Tobacco use in this population is unavoidable; therefore we determined years of tobacco use and tobacco dependency in CA and HC (see Supplementary Methods and Results). Psychiatric comorbidities, lifetime medication and polysubstance use are reported in Supplementary Table 3, 4 and 5 respectably. Participants were asked to abstain from crack cocaine as well as other drugs and alcohol, for at least 24 hours prior to the study and were urine-tested for the presence of the drugs before the MRI scan. The clinical and MRI sessions were done the same day as minimum and 4 days apart as maximum.

### MRI Acquisition

Brain images were acquired using a Phillips Ingenia 3T MR system (Phillips Healthcare, Best, Netherlands & Boston, MA, USA) with a 32-channel dS Head coil. We acquired structural T1-weighted data and DKI data using the fast kurtosis acquisition scheme. Specifically, the fast kurtosis protocol from Hansen *et al.*^30^ was employed requiring a total of 19 diffusion weighted volumes: 1 b = 0 scan for normalization and 9 distinct directions at each of b = 1000 s/mm2 and b = 2500 s/mm2. The data was acquired using a spin-echo single-shot EPI, TR/TE = 11820/115 ms, flip-angle = 90° and inversion recovery for CSF suppression. A total of 50 consecutive axial slices were acquired with isotropic resolution of 2 mm, matrix = 112 × 112, scan time = 4:46 min. T1-weighted images were acquired using a 3D FFE SENSE sequence, TR/TE = 7/3.5 ms, FOV = 240, matrix = 240 × 240 mm, 180 slices, gap = 0, plane = Sagittal, voxel 1 × 1 × 1 mm (5 participants were acquired with a voxel size =.75 ×.75 × 1 mm), scan time = 3.19 min. As part of a major addiction project, resting state fMRI and HARDI sequences were also acquired and are not part of this particular study. The order of the sequences was: rsfMRI, T1w, HARDI, fDKI, and was maintained across participants. Total scan time was around 50 min.

### T1-weighted preprocessing and image processing

T1-weighted images were converted from DICOM format to MINC for preprocessing. T1 images were preprocessed using an in-house preprocessing pipeline with the software Bpipe (http://cobralab.ca/software/mincbeast_bpipe.html) ^36^, which makes use of the MINC Tool-Kit (http://www.bic.mni.mcgill.ca/ServicesSoftware/ServicesSoftwareMincToolKit) and ANTs ^37^. Briefly, we performed N4 bias field correction ^38^, linear registration to MNI-space using ANTs, we cropped the region around the neck in order improve registration quality, followed by transformation back to native space.

The native space preprocessed files were input into the MAGeT-Brain morphological analysis pipeline (http://cobralab.ca/software/MAGeTbrain.html) ^39^. MAGeT-Brain is modified multi-atlas segmentation technique designed to take advantage of hard-to-define atlases and uses a minimal number of atlases for input into the segmentation process (atlases’ labels in Supplementary Methods and Results). We obtained segmentations and volumetric measures for whole striatum and thalamus, as well as their subdivisions. For shape analysis, indices of surface displacement ^20,40^ and surface area ^41^ for the striatum and thalamus were derived. Briefly, surface displacements were derived based on the average of the nonlinear portions of the 21 transformations estimated using MAGeT-Brain as the dot product between the surface normal and the local nonlinear registration vector at each point. Native surface area was estimated using a median surface representation based on the 21 surfaces from the MAGeT-Brain pipeline in native space. Surface area was estimated by assigning one-third of the surface area of each triangle to each vertex within the triangle. Finally, surface area and displacement values were blurred with a surface based diffusion-smoothing kernel of 5 mm ^42^. These measures were provided for the striatum (~6,300 vertices/hemisphere) and the thalamus (~3,000 vertices/hemisphere).

### DKI preprocessing

DKI data was corrected for motion and eddy currents using FSL version 5.0.9 (Oxford Centre for Functional MRI of the Brain). Tensor-based mean kurtosis (MKT) was calculated as previously described in Hansen *et al.* ^30^ using Matlab (MathWorks, Natick, Massachusetts). MKT was calculated on unsmoothed data as we aimed for ROI based statistics (it should be noted that motion and eddy current corrections introduce slight blurring due to resampling). T1-weighted images were skull-stripped ^43^ and linearly co-registered to the MKT images ^44^. To extract ROI based MKT values, we used the labels derived individually from the MAGeT Brain pipeline, focusing on whole left and right striatum, thalamus and nucleus accumbens (NAcc). The labels were resampled in DKI space using the T1 to DKI linear transformation, and mean MKT values from each label and each participant were calculated and stored in a CSV file for statistical analysis.

### Statistical Analysis

All data were analyzed using R Statistics and RStudio. All variables were tested for normality using the Shapiro-Wilks test. Participant data was analyzed using t-tests and chi-squared when appropriate. All tests were corrected for multiple comparisons using false discovery rate (FDR) ^45^ at 10% threshold and adjusted p-values are shown. Effect sizes were determined using sums of squares (SS) and partial-eta-squared (*partial-eta-squared = SS(effect) / (SS(effect) + SS(error))*. For striatum and thalamus whole volumes and subdivisions, we tested for group differences and group×age interactions using ANCOVA at an alpha level of 0.05 and using total brain volume, age (only group differences) and sex as covariates.

Also, we tested for linear relationships between years of consumption of crack cocaine in CA participants and striatum and thalamus volumes using multiple regression using total brain volume and sex as covariates with alpha of 0.05. *Post-hoc* we then performed two-tailed correlations between “years of consumption” and striatal and thalamic subnuclei volume. Finally, we performed a linear regression of the right pulvinar (rPul) volume and years consuming cocaine, with total brain volume, age and sex as covariates. A liberal FDR threshold was chosen due to the small sample size, the high variability of results in previous studies, and consistency with previous reports on shape analysis ^40,46^. For visualization purposes only, we created corrected volume variables using the residuals of the linear model.

Vertex-wise GLM analyses of surface morphometric measures were performed using RMINC (https://github.com/mcvaneede/RMINC), a statistical image analysis software package built to work in the R environment. All vertex-wise results were also corrected for multiple comparisons using FDR at 10% threshold. For surface area and displacement we tested for group differences and group×age interactions using GLM with total surface area, age and sex as covariates in surface area and only age and sex for displacement. We superimposed the striatum and thalamus subdivision atlases to label the subnuclei location of the significant clusters. Finally we created plots using the values of surface area and displacement observed in the peak vertex of significant clusters.

For the DKI data, we tested for group differences and group*×*age interactions (ANCOVA, α = 0.05) using age (only for the former) and sex as covariates. Again, FDR at 10% was used. We performed parametric correlation between years of consumption of crack cocaine (CA only) and striatum/thalamus MKT.

Finally, to better understand the results of the “age” interaction, we first performed two-tailed correlation between chronological age and years of consumption. Then we performed the ANCOVA analyses for striatum and thalamus volume, MKT and shape again using “years of consumption” as an additional covariate. We also performed multiple regression analysis of the thalamus MKT using “age” as a covariate to understand the influence of chronological age in the correlation.

## Results

Participant data are shown in Supplementary Table 1. There were no significant relationships between the morphological data and years of tobacco use in our data (see Supplementary Methods and Results for details on the analysis). Chronological age and years of consumption were positively correlated (r = 0.53, p = 0.001).

In whole volume we did not find group effects (HC > CA), though CA average volume was lower than HC in striatum and thalamus (Table 1). We found significant group×age interactions in striatum, in postcommisural caudate nucleus (PostCau), but not in the thalamus (Table 1). The interaction shows that striatum and caudate volume increased with age in the CA group and decreased in the HC (Figure 1). When “years of consumption” was included as a covariate, the group×age interactions in bilateral whole striatum were maintained (Table 2).

**Table 1.** Volumetric analysis.

| **Striatum** | **Volume (mean, sd)** |  | **Group Contrast** |  | **Group \* Age Interaction** |  |  |
| --- | --- | --- | --- | --- | --- | --- | --- |
|  | **HC** | **CA** | **F (1,71)** | **p** | **F (1,70)** | **p** | **pFDR** |
| lStria | 9251 (±1011) | 9114 (±708) | 1.02 | 0.32 | 4.31 | 0.04 | 0.08* |
| rStria | 9248 (±1003) | 9130 (±679) | 0.74 | 0.39 | 5.18 | 0.02 | 0.08* |
| lPrePut | 1241 (±192) | 1204 (±166) | 1.07 | 0.31 | 0.87 | 0.35 | 0.75 |
| rPrePut | 1295 (±185) | 1261 (±159) | 1.04 | 0.31 | 0.59 | 0.44 | 0.75 |
| lPosPut | 3867 (±432) | 3802 (±337) | 0.94 | 0.34 | 1.14 | 0.29 | 0.75 |
| rPosPut | 3606 (±418) | 3552 (±321) | 0.70 | 0.41 | 1.07 | 0.30 | 0.75 |
| lPreCau | 2149 (±277) | 2121 (±212) | 0.38 | 0.54 | 4.59 | 0.04 | 0.38 |
| rPreCau | 2269 (±261) | 2253 (±258) | 0.10 | 0.76 | 3.39 | 0.07 | 0.56 |
| lPosCau | 1389 (±186) | 1396 (±140) | 0.04 | 0.84 | 6.04 | 0.02 | 0.27 |
| rPosCau | 1406 (±206) | 1429 (±187) | 0.39 | 0.53 | 10.25 | 0.002 | 0.06* |
| lVS | 1152 (±146) | 1130 (±109) | 0.88 | 0.35 | 1.23 | 0.27 | 0.75 |
| rVS | 1163 (±130) | 1151 (±119) | 0.28 | 0.60 | 2.58 | 0.11 | 0.72 |
| **Thalamus** | **HC** | **CA** | **F (1,71)** | **p** | **F (1,70)** | **p** | **pFDR** |
| lThal | 5752 ± 503 | 5641 ± 512 | 1.47 | 0.23 | 0.03 | 0.87 | 0.97 |
| rThal | 6124 ± 506 | 6024 ± 509 | 1.31 | 0.26 | 0.53 | 0.47 | 0.49 |
| lLGN | 134 (±20) | 135 (±18) | 0.01 | 0.93 | 0.01 | 0.94 | 0.99 |
| rLGN | 213 (±29) | 214 (±29) | 0.01 | 0.93 | 0.001 | 0.98 | 0.99 |
| lMGN | 178 (±21) | 175 (±20) | 0.35 | 0.56 | 1.51 | 0.22 | 0.75 |
| rMGN | 204 (±26) | 103 (±20) | 0.06 | 0.80 | 0.01 | 0.94 | 0.99 |
| lAN | 111 (±22) | 107 (±17) | 0.99 | 0.32 | 0.50 | 0.48 | 0.75 |
| rAN | 163 (±26) | 156 (±28) | 1.60 | 0.21 | 2.27 | 0.14 | 0.75 |
| lCN | 290 (±29) | 284 (±27) | 1.22 | 0.27 | 0.53 | 0.47 | 0.75 |
| rCN | 192 (±20) | 190 (±18) | 0.47 | 0.50 | 0.60 | 0.44 | 0.75 |
| lLD | 53 (±9) | 54 (±10) | 0.21 | 0.65 | 0.06 | 0.81 | 0.99 |
| rLD | 60 (±8) | 58 (±9) | 1.02 | 0.32 | 0.06 | 0.94 | 0.99 |
| lLP | 421 (±64) | 422 (±58) | 0.01 | 0.93 | 0.97 | 0.33 | 0.75 |
| rLP | 319 (±44) | 317 (±48) | 0.02 | 0.90 | 0.79 | 0.38 | 0.75 |
| lMD | 952 (±99) | 910 (±121) | 2.92 | 0.09 | 0.42 | 0.52 | 0.75 |
| rMD | 1050 (±107) | 1020 (±121) | 1.48 | 0.23 | 0.93 | 0.34 | 0.75 |
| lPul | 1571 (±196) | 1564 (±193) | 0.03 | 0.87 | 1.65 | 0.28 | 0.75 |
| rPul | 2005 (±243) | 1994 (±218) | 0.07 | 0.80 | 1.65 | 0.20 | 0.75 |
| lVAN | 433 (±53) | 421 (±56) | 1.28 | 0.26 | 0.005 | 0.94 | 0.99 |
| rVAN | 545 (±63) | 535 (±93) | 0.21 | 0.65 | 0 | 0.99 | 0.99 |
| lVLN | 803 (±89) | 778 (±101) | 1.90 | 0.17 | 0.49 | 0.49 | 0.75 |
| rVLN | 662 (±62) | 648 (±89) | 0.69 | 0.41 | 0.06 | 0.81 | 0.99 |
| lVPN | 553 (±64) | 548 (±62) | 0.19 | 0.66 | 0.16 | 0.69 | 0.99 |
| rVPN | 507 (±62) | 498 (±55) | 0.55 | 0.46 | 0.12 | 0.73 | 0.97 |

Volume is in mm<sup>3</sup>, sd = standard deviation, HC = Healthy controls, CA = Cocaine addicts, p = uncorrected p, pFDR = adjusted p at FDR 10%, l = left, r = right, Stria = whole striatum, Thal = whole thalamus, Pre = precommissural, Pos = postcommissural, Put = putamen, Cau = caudate, VS = ventral striatum, LGN = left lateral geniculate nucleus, MGN = medial geniculate nucleus, AN = anterior nuclei, CN = central nuclei, LD = lateral dorsal, LP = lateral posterior, MD = medial dorsal, Pul = pulvinar, VAN = ventral anterior nucleus, VLN = ventral lateral nucleus, VPN = ventral posterior nucleus, \* pFDR < 0.10.

There were no significant relationships between years consuming crack cocaine and whole striatum/thalamus volume. The post-hoc correlations with the subnuclei found no correlation with years consuming cocaine in the striatum. In the thalamic subdivisions, we found a significant correlation between years consuming with right pulvinar volume (r= -.52, uncorrp = 0.001, pFDR = 0.05). The multiple regression analysis confirmed this relationship (Supplementary Table 7). Thus, the longer years of crack cocaine consumption the smaller right pulvinar volume. Introducing “age” as a covariate to the multiple regression model did not affect the R^2^, but it did reduce the significance and beta value of years of consumption (Table 2).

**Table 2.**
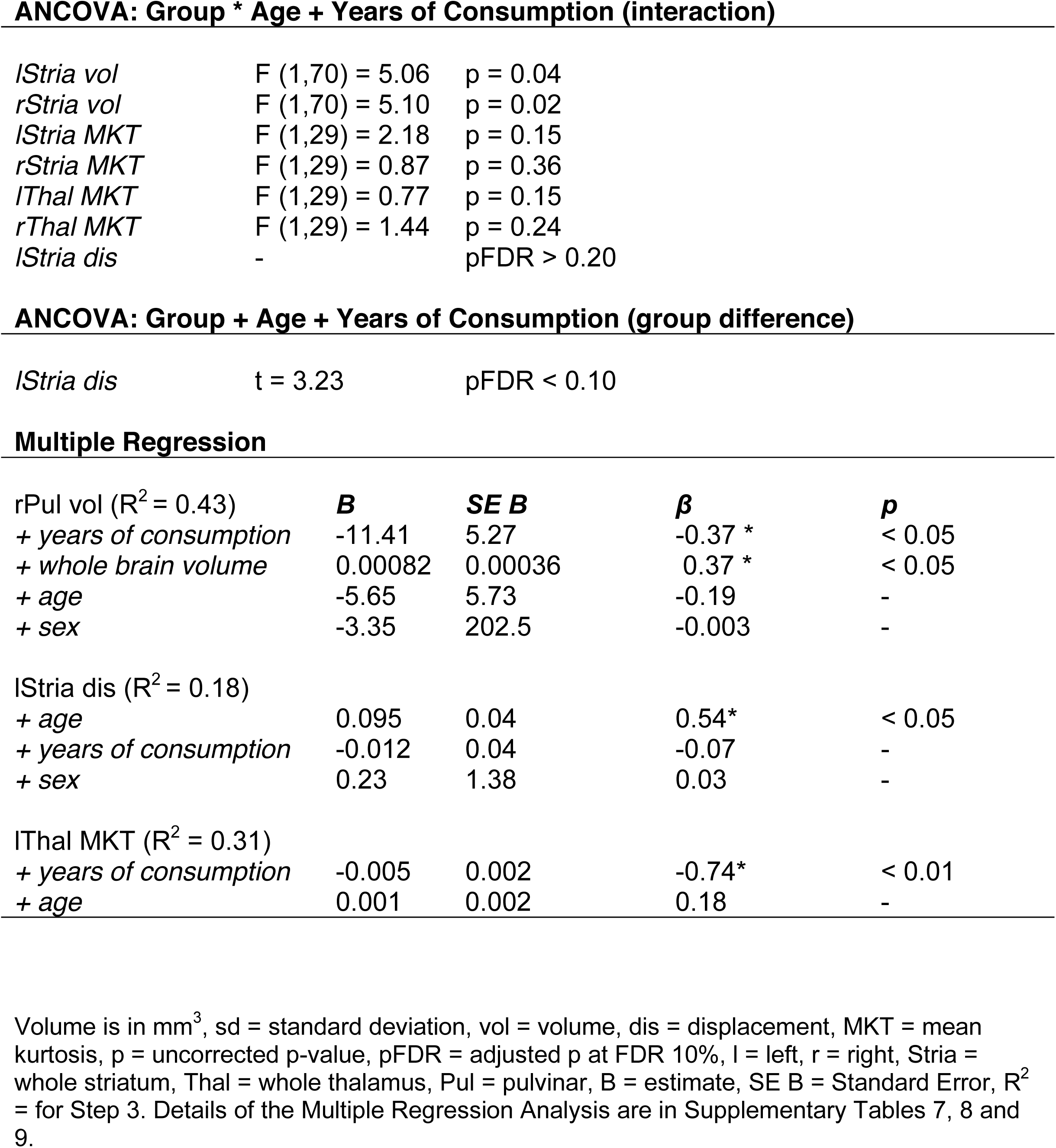
Years of consumption and age covariates analysis.

|  |  |  |
| --- | --- | --- |
| *lStria vol* | F (1,70) = 5.06 | p = 0.04 |
| *rStria vol* | F (1,70) = 5.10 | p = 0.02 |
| *lStria MKT* | F (1,29) = 2.18 | p = 0.15 |
| *rStria MKT* | F (1,29) = 0.87 | p = 0.36 |
| *lThal MKT* | F (1,29) = 0.77 | p = 0.15 |
| *rThal MKT* | F (1,29) = 1.44 | p = 0.24 |
| *lStria dis* | - | pFDR > 0.20 |

**ANCOVA: Group + Age + Years of Consumption (group difference)**

|  |  |  |
| --- | --- | --- |
| *lStria dis* | t = 3.23 | pFDR < 0.10 |

**Multiple Regression**
|  |  |  |  |  |
| --- | --- | --- | --- | --- |
| *rPul vol* (R<sup>2</sup> = 0.43) | **B** | **SE B** | **$\beta$** | **p** |
| + years of consumption | -11.41 | 5.27 | -0.37 * | < 0.05 |
| + whole brain volume | 0.00082 | 0.00036 | 0.37 * | < 0.05 |
| + age | -5.65 | 5.73 | -0.19 | - |
| + sex | -3.35 | 202.5 | -0.003 | - |
| *lStria dis* (R<sup>2</sup> = 0.18) |  |  |  |  |
| + age | 0.095 | 0.04 | 0.54* | < 0.05 |
| + years of consumption | -0.012 | 0.04 | -0.07 | - |
| + sex | 0.23 | 1.38 | 0.03 | - |
| *lThal MKT* (R<sup>2</sup> = 0.31) |  |  |  |  |
| + years of consumption | -0.005 | 0.002 | -0.74* | < 0.01 |
| + age | 0.001 | 0.002 | 0.18 | - |

Volume is in mm<sup>3</sup>, sd = standard deviation, vol = volume, dis = displacement, MKT = mean kurtosis, p = uncorrected p-value, pFDR = adjusted p at FDR 10%, l = left, r = right, Stria = whole striatum, Thal = whole thalamus, Pul = pulvinar, B = estimate, SE B = Standard Error, R<sup>2</sup> = for Step 3. Details of the Multiple Regression Analysis are in Supplementary Tables 7, 8 and 9.

**Figure 1.**
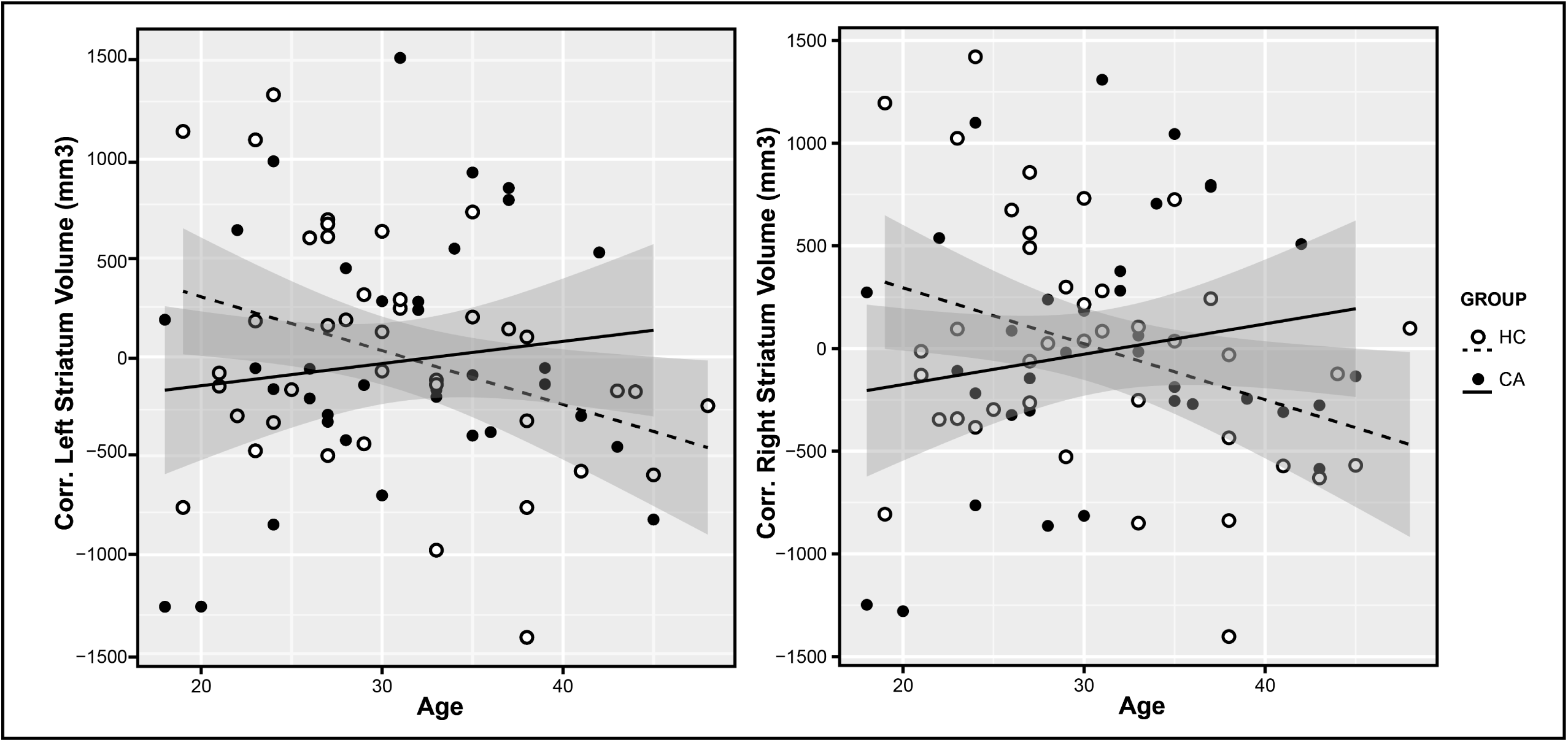
Striatum volume group×age interaction. Scatter plot of left and right striatum corrected volume with regression lines and shadowed confidence intervals at 0.95 per group. Group×Age interaction statistics, left striatum: F (1,70) = 4.31, uncorrp = 0.04, pFDR = 0.08, es = 0.06; right striatum: F (1,70) = 5.18, uncorrp = 0.02, pFDR = 0.08, es = 0.05. HC = healthy controls, CA = cocaine addicts.

In displacement, we found a significant group difference (HC > CA) in the left striatum (t = 3.14, pFDR < 0.10) localized in the ventral striatum/NAcc area (Figure 2). We also found a non-significant group×age interaction (t = 3.28, pFDR < 0.20) in the same area. In general, the CA group had more contraction (or inward displacement) than HC in the cluster peak. Introducing the “years of consumption” covariate for the left striatum displacement, we found a consistent significant group difference with slightly increased t-value, although the group×age interaction became weaker (Table 2). The left striatum displacement cluster peak vertex was not significantly correlated with years consuming (r= 0.23, p = 0.19), however it was positively correlated with age (r = 0.48, p = 0.003). The multiple regression analysis of the cluster peak showed an influence solely by age (Table 2 and Supplementary Table 8). There were no significant clusters in the thalamus. The surface area did not show significant clusters in striatum and thalamus.

**Figure 2.**
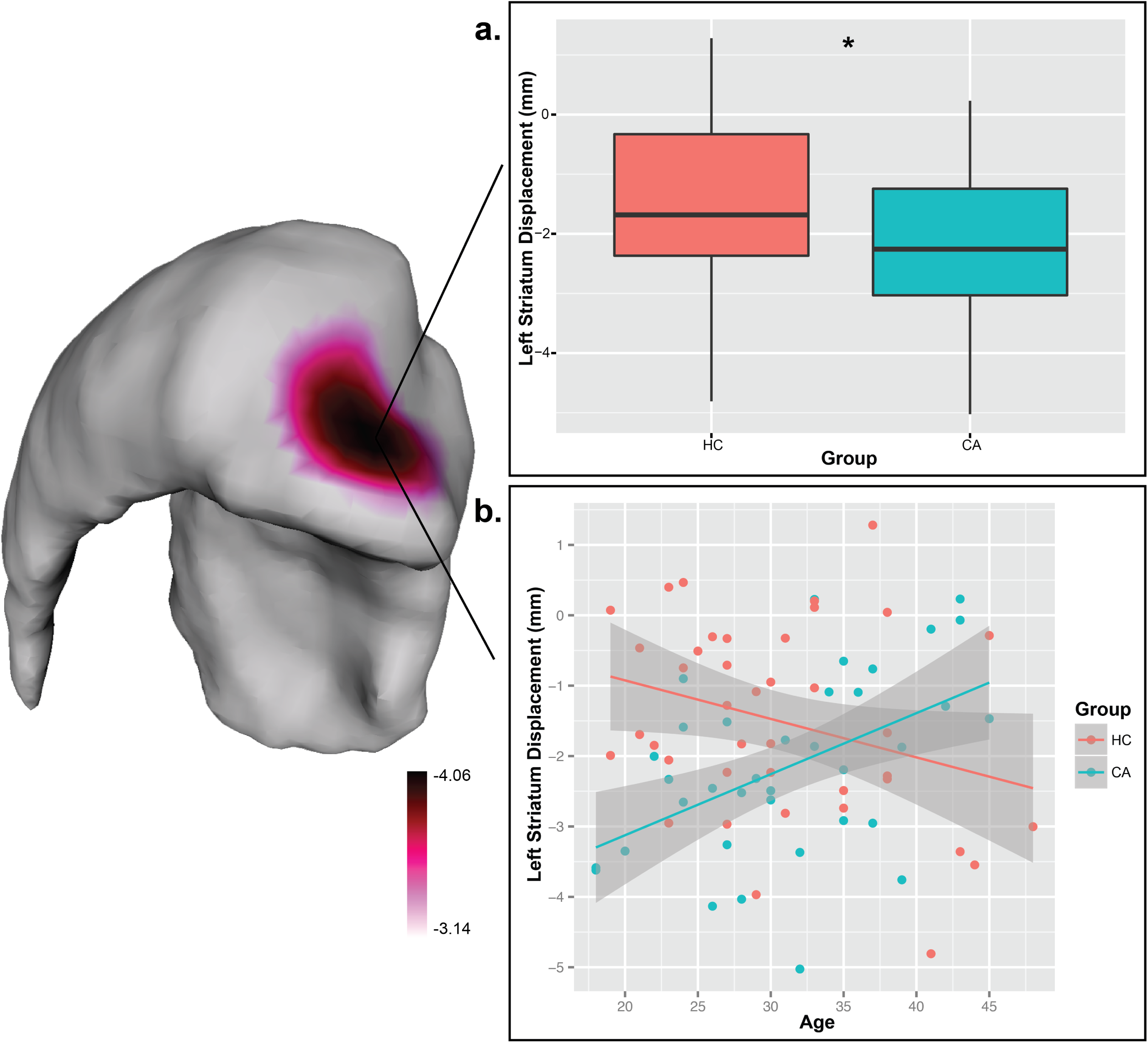
Striatum displacement. Left: Caudal anterior view of the left striatum showing the significant group cluster on ventral striatum/NAcc. Right, a) boxplot of the significant (*) displacement difference between groups in the peak vertex location (pFDR 10%) t = 3.14, pFDR < .10; b) scatter plot of the group*age interaction in displacement showing regression lines and shadowed confidence intervals at 0.95 per group. The color bar shows the t-values at pFDR threshold of 20% (t = 3.28, pFDR < .20). Negative values = contraction, positive values = expansion, HC = healthy controls, CA = cocaine addicts.

The demographics of the fDKI smaller sample size were similar to the volume analysis. The fDKI analysis did not show a group effect. However, it showed significant group×age interactions in bilateral striatum (left: F (1, 30) = 4.49, uncorrp = 0.04, pFDR = 0.05, es = 0.13; right: (F (1, 30) = 3.81, uncorrp = 0.06, pFDR = 0.06, es = 0.11), and in bilateral thalamus (left: F (1, 30) = 6.12, uncorrp = 0.02, pFDR = 0.05, es = 0.17; right: F (1, 30) = 5.57, uncorrp = 0.03, pFDR = 0.05, es = 0.16), with a higher effect size on left thalamus (Figure 3). Mostly in the thalamus, MKT decreased with age in CA while it increased in HC, though it did show a different pattern in striatum. When “years of consumption” was included as a covariate, the group×age interactions in bilateral striatum and thalamus were not maintained (Table 2).

Striatum MKT showed no significant correlations. We found significant negative correlations between years consuming with left thalamus MKT (r= -0.62, uncorrp = 0.008, pFDR = 0.03) and not significant with right thalamus MKT (r= -0.47, uncorrp = 0.06, pFDR = 0.1), meaning the longer years they consumed the lower MKT values. The multiple regression analysis with age as covariate maintained the significant correlation in left thalamus MKT (Table 2 and Supplementary Table 9). We did not find significant correlations between striatum and thalamus MKT and volume.

**Figure 3.**
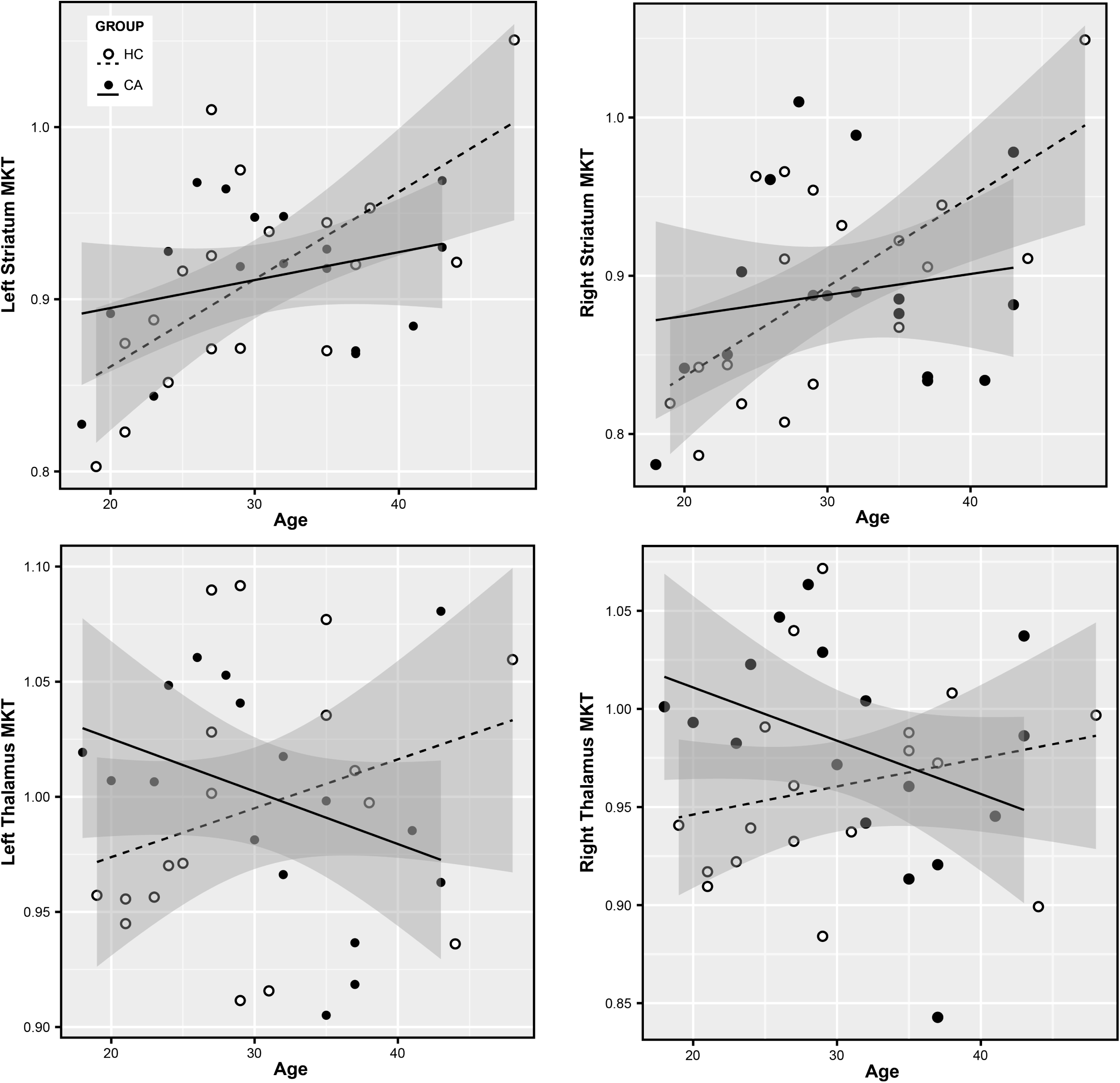
Striatum and thalamus MKT group×age interaction. Scatter plot of (top) striatum MKT and (bottom) thalamus MKT with regression lines and shadowed confidence intervals at 0.95 per group. Group×Age interactions, left striatum: F (1, 30) = 4.49, uncorrp = 0.04, pFDR = 0.05, es = 0.13; right striatum: F (1, 30) = 3.81, uncorrp = 0.06, pFDR = 0.06, es = 0.11; left thalamus: F (1, 30) = 6.12, uncorrp = 0.02, pFDR = 0.05, es = 0.17; right thalamus: F (1, 30) = 5.57, uncorrp = 0.03, pFDR = 0.05, es = 0.16. HC = healthy controls, CA = cocaine addicts, MKT = mean kurtosis.

## Discussion

Our findings suggest that morphometric changes in active crack cocaine addiction are subtle, and that volumetric analysis is less sensitive than shape analysis and DKI to measure these changes. However, we also found that volume and shape in the striatum were more related to abnormal aging in cocaine addicts, while morphometric differences in thalamus and DKI differences in both striatum and thalamus were more related to years consuming the drug. In our sample, striatum and thalamus volume was not different between healthy controls and crack cocaine addicts. This agrees with the study by Martinez *et al* ^11^ where they also showed no significant volumetric differences in striatum using manual segmentation. In their study, they also admitted patients with active crack cocaine addiction and had a smaller sample size than ours. Narayana *et al.* ^12^ did not find differences as well using whole-brain voxel-based morphometry (VBM) with a similar sample size, without specifying the type of cocaine delivery. In studies that include the thalamus, polysubstance users had greater volume in the right thalamus compared to cocaine-users, and another study found that cocaine-dependent subjects have decreased volume in the left thalamus compared to healthy controls ^10^. Inability to detect group differences could be the result of several factors: variability of cocaine consumption, quality of the cocaine used, polysubstance use, or type of healthy controls (HC) used for comparison. In our case, the HC were closely paired for age, sex, handedness and education. We found that striatum volume increased with age in our CA group, and the effect was more prominent in the left postcommissural caudate. In healthy human population, striatum and thalamus volumes decrease with normal aging, suggesting age-related atrophy in our sample ^47,48^. Bartzokis *et al.*^49^ showed accelerated age-related decline in cortical volume in cocaine addicts, though they did not study striatum nor thalamus specifically. Gray matter volume decrease shown by MRI is related to a decrease in synaptic spine density in murine models ^50^. The volume increase in our CA group could suggest an increase in dendritic spines, supporting changes in synaptic plasticity due to years of cocaine consumption ^34^. However, volume could be affected by non-specific microstructural changes; hence, an increase in volume by itself does not necessarily suggest an increase in dendritic spines. Instead, it could also suggest reorganization in circuitry, scarring or inflammation ^34,51,52^. Our analysis also showed that years of consumption did not affect volume in the striatum, which seems to imply that the differences found are related to abnormal aging. However, we found a positive medium-high correlation between years of consumption and age, which suggests these variables are difficult to differentiate and considered in the interpretation.

Thalamus volume was not affected by age in the CA group as in striatum. A deeper investigation showed that longer years consuming crack cocaine were associated with decreased right pulvinar volume. Age did not affect the statistical model, therefore we suspect the effect may be more related to years of consumption. The pulvinar is a thalamic nucleus highly connected to the visual cortex and the superior colliculus, and recent studies suggest one of its main functions is visual attention and modulation ^53^. It is highly connected to the striatum as well ^54^ and related to the addictive pathology ^34^. In fact, striatal availability of dopamine D2 receptor predicts thalamic functional responses to reward in cocaine addicts ^55^. This suggests cocaine abuse may be related to atrophy of the thalamus and could be part of the habit reinforcement stage of the addiction cycle theory^56^. If the volume decrease is related to a decrease in dendritic spines, it may also explain inhibition and attention issues seen in cocaine addiction due to the visual-motor connectivity of the pulvinar ^51^. However, at the moment this is only possible to investigate in murine models.

The shape analysis showed significant changes in striatum anatomy between the groups, as opposed to volume. We found a greater contraction (or inward displacement) in the CA group than HC in the ventral striatum/NAcc, an important part of the reward system and addiction. The NAcc seems to be involved in the intoxication and withdrawal stages of cocaine addiction, with the consequent drug seeking behavior ^34,56^. A study in adults with prenatal exposure to cocaine also found displacement differences compared to healthy controls in striatum using shape analysis ^57^. However, they only studied the caudate and putamen, and their results were not corrected for multiple comparisons, though they support our own findings. A murine model of adolescent cocaine exposure determined displacement differences in striatum, with significant expansion of the lateral surface and contraction of the medial surface after 30 days of abstinence ^3^. We also found a non-significant age-related shape interaction with group, showing expansion of the ventral striatum/NAcc in the CA group. This is similar to the findings in volume. When we included years of consumption in the statistical model as another covariate, the group difference was maintained but the group×age interaction disappeared, suggesting shape may represent changes related to age and not years of consumption. A study on schizophrenia suggests that the ventral striatum/NAcc contracts with age in healthy controls ^20,41^, and this seems to be true for thalamus as well ^58^. Our HC group showed the same age-related contraction, while the CA group showed expansion, though this was not significant. This evidence may suggest that, although the NAcc is contracted in crack cocaine addiction, there is a tendency to expand with age. Similarly to volume, this could be related to either an increase of dendritic spines or gliosis. To our knowledge, this is the first shape analysis study in cocaine-addicted humans; therefore bigger sample sizes and longitudinal studies are needed to corroborate these results. It is relevant to mention that the MAGeT algorithm we applied in this study, has been shown to give similar results to the gold standard (manual segmentation; achieving Dice Kappa against manual segmentation of ~ 0.88 – 0.89), as opposed to other tools that over or under estimate the volumes ^39^.

The DKI analysis, as with the volume, did not show group differences in striatum and thalamus MKT. It did show an age-related group interaction in striatum and thalamus with low effect sizes. MKT decreased with age in the CA group, especially evident in thalamus, and increased with age in HC. We compare our results with the recently introduced MKT directly to studies employing conventional MK. Falangola *et al.*^28^ found whole-brain MK to increase rapidly with age until age 18, followed by a less steep increase until age 47 after which a decline in MK was observed. For gray matter (GM) the MK was seen to continue to increase through life, whereas for white matter (WM) MK was found to increase rapidly up to age 18 and then steadily decline. In agreement with that study, MKT increased with age in HC, but decreased with aging in the CA group, especially evident in thalamus. Meaning that in our study, crack cocaine addicts show an abnormal MKT decline with age, as well as decrease in thalamic MKT with years of cocaine use. Interestingly, introducing years of consumption negatively affected the model, and we found significant correlation between thalamic MKT and years of consumption. This may suggest that MKT may reflect both substance-related and age-related alterations, with a higher specificity for the former. A decreased MKT compared to normal brain tissue would suggest a less complex tissue microstructure with fewer diffusion barriers causing the diffusion process to be less non-Gaussian. Specific causes for this could be loss of neurites, leading to fewer cell connections, decreased tissue integrity and increased extra-cellular space. This may be corroborated by the significant negative correlations with years of cocaine use that, when compared with the pulvinar volume correlation, could suggest a reflection of thalamic atrophy. To our knowledge, this is the first study to use diffusion kurtosis imaging in substance addiction, hence more studies are needed in this and other types of substance abuse.

There were several limitations to our study. With the low effect sizes in all the morphological contrasts, it seems higher samples sizes may be needed to find more significant differences in these areas. Nevertheless, we were able to obtain significant results with our limited sample sizes. The DKI analysis had the lowest sample size, and still it showed a similar effect size than the other analyses. And unlike the rest of the analyses, the DKI showed thalamic effects as well as striatal. Our significant threshold for the multiple comparisons FDR was 10% (q = 0.1), which may be considered more liberal than usual and caution should be taken when interpreting our findings. However, this approach was preferred to allow for a more exploratory study and we have successfully used it in our previous studies. With our sample size (n = 76) and with the largest effect size in volume (group×age interaction of postcommisural caudate nucleus, es = 0.13), we achieved a power of 90%. However, with the lowest significant effect size in volume (group×age interaction of striatum, es = 0.05), we achieved a power of 50%. To achieve an 80% power at an alpha = 0.05, with those effect sizes, we would need a sample size of at least n = 152. This was calculated using the software G*Power (http://www.gpower.hhu.de/en.html) for an ANCOVA with Fixed effects, main effects and interactions, 2 groups and 3 covariates. Tobacco use and dependency was prevalent in half of our CA group, and although we did not find any effects in striatum and thalamus, one study has shown striatal volume and shape relation to craving tobacco ^40^. Finally, age and years of consumption were positively correlated, and this colinearity should be considered when interpreting the results. One thing to take into consideration is that the variable “years of consumption” may be somewhat subjective, as it is investigated by the clinician but reported by the patient and sometimes by family members. Studies in murine models may help to model the nature of the effects of poly-substance abuse, age and years of consumption ^3^.

Our findings show morphological and microstructural changes in crack cocaine addicts, related to age and years of cocaine consumption. Most importantly, we found that striatal morphological changes may be more related to age, while thalamic changes may be more related to years consuming. We also found abnormal MKT development with age and years of consumption, highlighting the potential for MKT, and DKI in general, as a method for in vivo investigation of the effects of addiction on brain microstructure. The shape analysis showed contraction of the nucleus accumbens when compared to healthy controls. We showed that volumetric analysis by itself provides incomplete information about morphology and suggest that shape analysis and/or diffusion kurtosis imaging should be used in addition to better characterize brain pathology in substance abuse.

### Funding and Disclosure

This project was funded by CONACYT-FOSISS project No. 0201493 and CONACYT-Cátedras project No. 2358948. The authors declare no conflict of interest.

## Acknowledgements

We would like to thank the people who helped this project in one way or another: Francisco J. Pellicer Graham, Margarita López-Titla, Aline Leduc, Erik Morelos-Santana, Diego Angeles, Alely Valencia, Laura Arreola, Lya Paas, Daniela Casillas, Sarael Alcauter, Luis Concha and Bernd Foerster. We would also like to thank Rocio Estrada Ordoñez and Isabel Lizarindari Espinosa Luna at the Unidad de Atención Toxicologica Xochimilco for all their help and effort. Finally, we would like to thank the study participants for their cooperation and patience.

## Supplementary Methods and Results

### Methods

#### Participants

The diagnostic interview for participants was a clinical interview, the MINI-Plus, and the Structured Clinical Interview for DSM Axis II (SCID-II), by a trained psychiatrist. To confirm substance use, we did the urine test “instant-view” which detects amphetamines, methamphetamines, benzodiazepines, cocaine, cannabis and opioids. This was made just before or after obtaining the MRI to know if there were any drugs in the organism. To study the lifetime use of other substances we used the Addiction Severity Index (ASI) (see Supplementary Table 5). Psychiatric comorbidities are shown in Supplementary Table 3 and prescribed medication in Supplementary Table 4.

Tobacco dependency was based on the “nicotine/tobacco dependency” description in the DSM-IV by answering three questions: 1) Do you smoke more than five cigarettes per day? 2) Have you tried to quit smoking, but failed? 3) If you stop smoking, do you feel a physical discomfort? An answer of “yes” to all three questions was reported as positive “tobacco dependency”. “Years of tobacco use” was a dependent variable in a preliminary analysis. We tested the relationship between years of tobacco use and the morphological data before the general analysis to avoid confounds with striatum as it has been shown that tobacco craving is related to striatum morphology (Janes et al, 2015).

### Clinical and Cognitive Instruments

As part of the main ongoing addiction project where this study is part of, there were several clinical standardized questionnaires and cognitive tests applied to the participants. Trained psychiatrists and psychologists working on the project applied the clinical tests, questionnaires and interviews. A trained psychology student applied the cognitive tests in a dedicated room using a computer (except those in written form). These were not used for our study. The different questionnaires and tests were as follows:

Clinical (See Supplementary Table 2)

Cognitive Tests

- Digit Span
- Letter & Numbers
- Brown-Peterson procedure
- Wisconsin Card Sorting Test
- Flanker Task
- Go-No Go Task
- Tower of London
- Iowa Gambling Test
- Big Five
- Cognitive Estimation Test
- Baron-Cohen Reading the Mind in the Eyes

For MAGeT-Brain, in this case, we use an atlas of the basal ganglia and thalamus previously derived from the reconstruction of serial histological data and warped to fit an MRI model (Chakravarty et al, 2006). MAGeT-Brain bootstraps the segmentation through the population under study by performing atlas-based segmentation on a subset of 21 individuals (11 HC/10 CA). We then use these templates to provide 21 candidate labels for each subject using nonlinear registration via the ANTs algorithm (Avants et al, 2011); labels are then fused using majority-vote (retaining the most frequently occurring label at each voxel). Each segmentation was verified using a quality control image.

### Structures studied

The atlases in this folder are based on the Colin-27 Average Brain (http://www.bic.mni.mcgill.ca/ServicesAtlases/Colin27) (Chakravarty et al. 2006)

-Whole thalamus, Globus Pallidus and Striatum

Label: 1 - left striatum
Label: 2 - left globus pallidus
Label: 3 - left thalamus
Label: 4 - right striatum
Label: 5 - right globus pallidus
Label: 6 - right thalamus

-Subdivisions of the striatum

Label: 100 - left nucleus accumbens / ventral striatum
Label: 101 - left precommissural putamen
Label: 102 - left precommissural caudate
Label: 104 - left postcommissural caudate
Label: 105 - left postcommissural putamen
Label: 110 - right nucleus accumbens / ventral striatum
Label: 111 - right precommissural putamen
Label: 112 - right precommissural caudate
Label: 114 - right postcommissural caudate
Label: 115 - right postcommissural putamen

-Subdivisions of the thalamus

Label: 1 - left lateral geniculate nucleus (LGN)
Label: 2 - left medial geniculate nucleus (MGN)
Label: 3 - left anterior nuclei
Label: 4 - left central nuclei
Label: 5 - left lateral dorsal
Label: 6 - left lateral posterior
Label: 7- left medial dorsal
Label: 8 - left pulvinar
Label: 9 - left ventral anterior nucleus
Label: 10 - left ventral lateral nucleus
Label: 11 - left ventral posterior nucleus
Label: 21 - right lateral geniculate nucleus (LGN)
Label: 22 - right medial geniculate nucleus (MGN)
Label: 23 - right anterior nuclei
Label: 24 - right central nuclei
Label: 25 - right lateral dorsal
Label: 26 - right lateral posterior
Label: 27 - right medial dorsal
Label: 28 - right pulvinar
Label: 29 - right ventral anterior nucleus
Label: 30 - right ventral lateral nucleus
Label: 31 - right ventral posterior nucleus

To understand the relationship between the MKT and the morphological variables, we performed correlations between left and right striatum/thalamus MKT and volume, and then performed a GLM analysis of only thalamus surface area and MKT. Striatum was not tested because no MKT/volume correlation was found here.

## Results

We found significant regression between surface area and MKT in the left thalamus (t= 3.82, pFDR = 0.05), specifically in the area of the left ventral posterior nucleus, and in right thalamus (t= 3.21, pFDR = 0.1) in the area of the right anterior nuclei and right ventral posterior nucleus (Supplementary Figure 2).

---

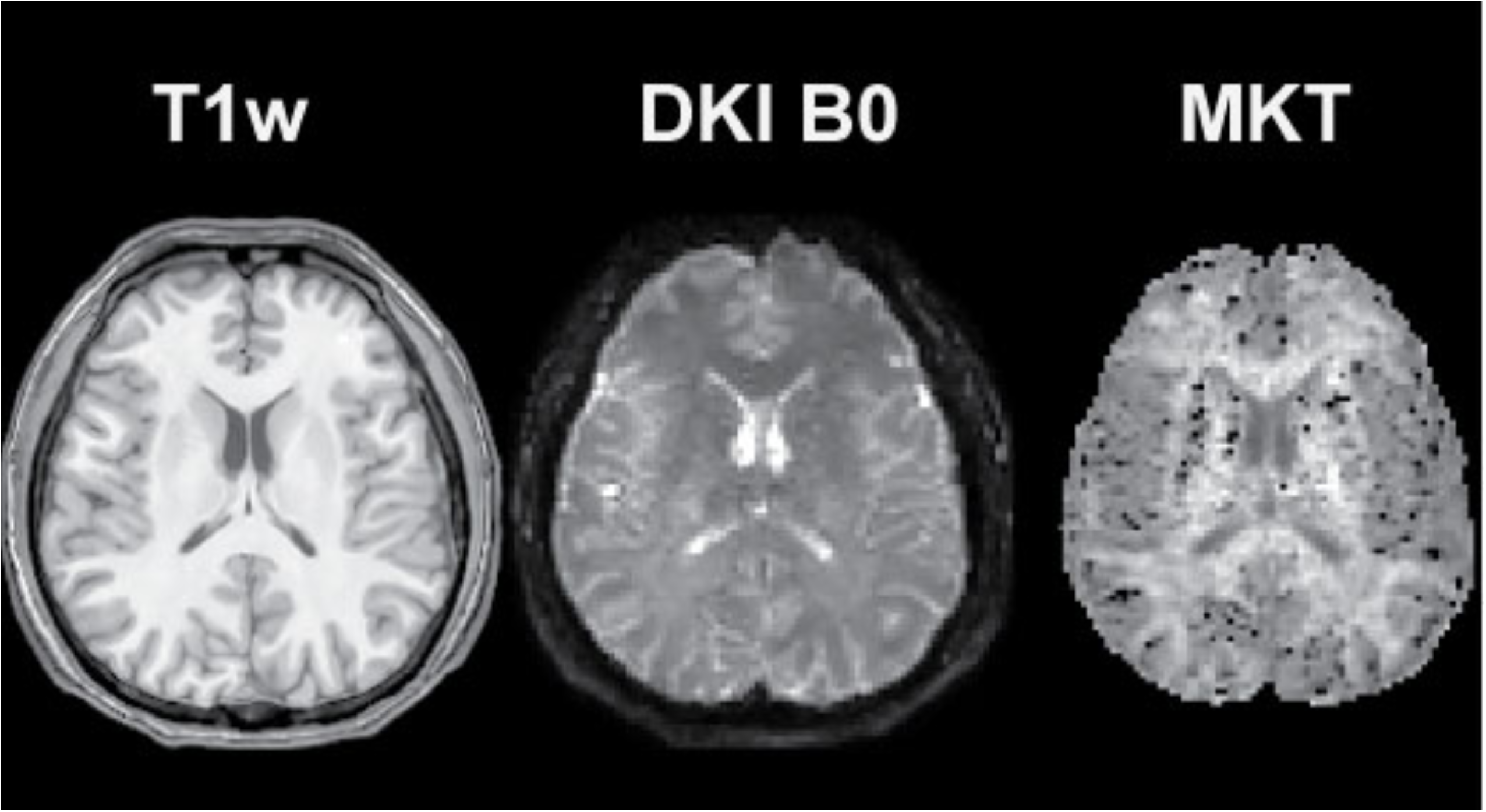

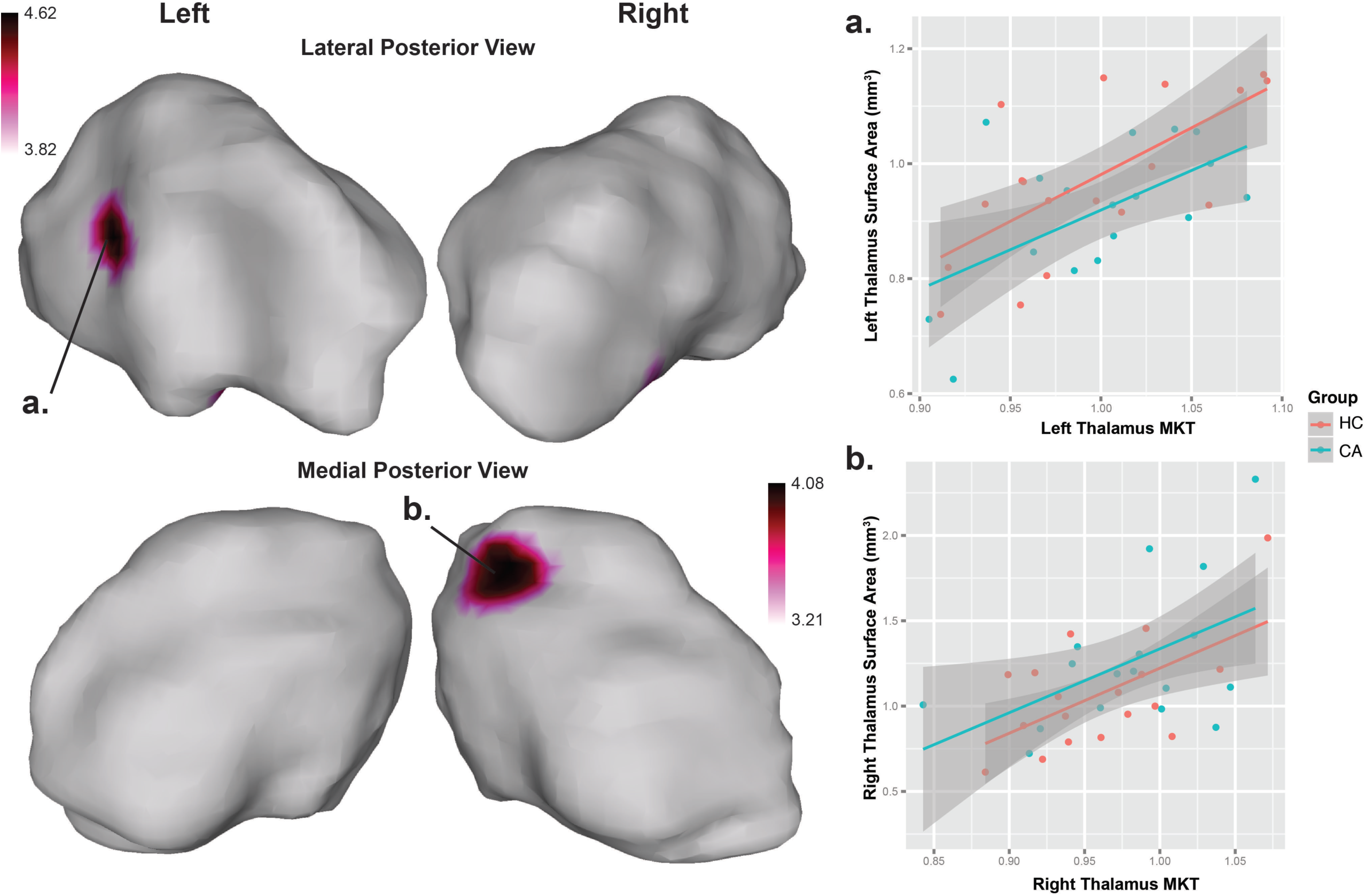

**Supplementary Table 1.** Participant data.

|  | CA (n=36) | HC (n=40) | Stats |
| --- | --- | --- | --- |
| Age | 31.5 (18 – 45) | 29.5 (19 – 48) | $t = -0.43$, $p = 0.67$ |
| Education | Secondary | High school | $\chi^2 = 13.87$, $p = 0.02$ |
| Handedness ** | (n = 36) | (n = 38) | $\chi^2 = 0.07$, $p = 0.96$ |
| Right | 30 | 31 |  |
| Left | 4 | 5 |  |
| Ambidextrous | 2 | 2 |  |
| Onset age of consumption | (n = 35), 20 (13 – 41) | na | - |
| Years consuming | (n = 34), 9 (1 – 25) | na | - |
| Average consumption per intake ** | (n = 33) | na | - |
| .8 gm | 0 | na | - |
| 1.6 – 2.4 gm | 1 | na | - |
| 3.2 – 5.6 gm | 6 | na | - |
| 8 – 9 gm | 16 | na | - |
| 9 – 10 gm | 9 | na | - |
| > 10 gm | 1 | na | - |
| Monthly expense in cocaine (USD)* | (n = 38), \$145 (± \$180) | na | - |
| Tobacco dependency ** | (n = 33), 19 | (n = 40), 3 | $\chi^2 = 19.22$, $p < .001$ |
| Years of tobacco use | (n = 32), 14.03 (±8.81) | (n = 40), 5.88 (±7.94) | $t = -4.08$, $p < 0.001$ |

Median (min – max) for all except: \* = mean (sd), \*\* = count. CA = Crack cocaine addicts, HC = Healthy controls, n = due to absence of data we show each sample size per variable, na = not applicable, gm = grams, Stats = statistics, t = t-value, $\chi^2$ = chi-squared.

**Supplementary Table 2.**
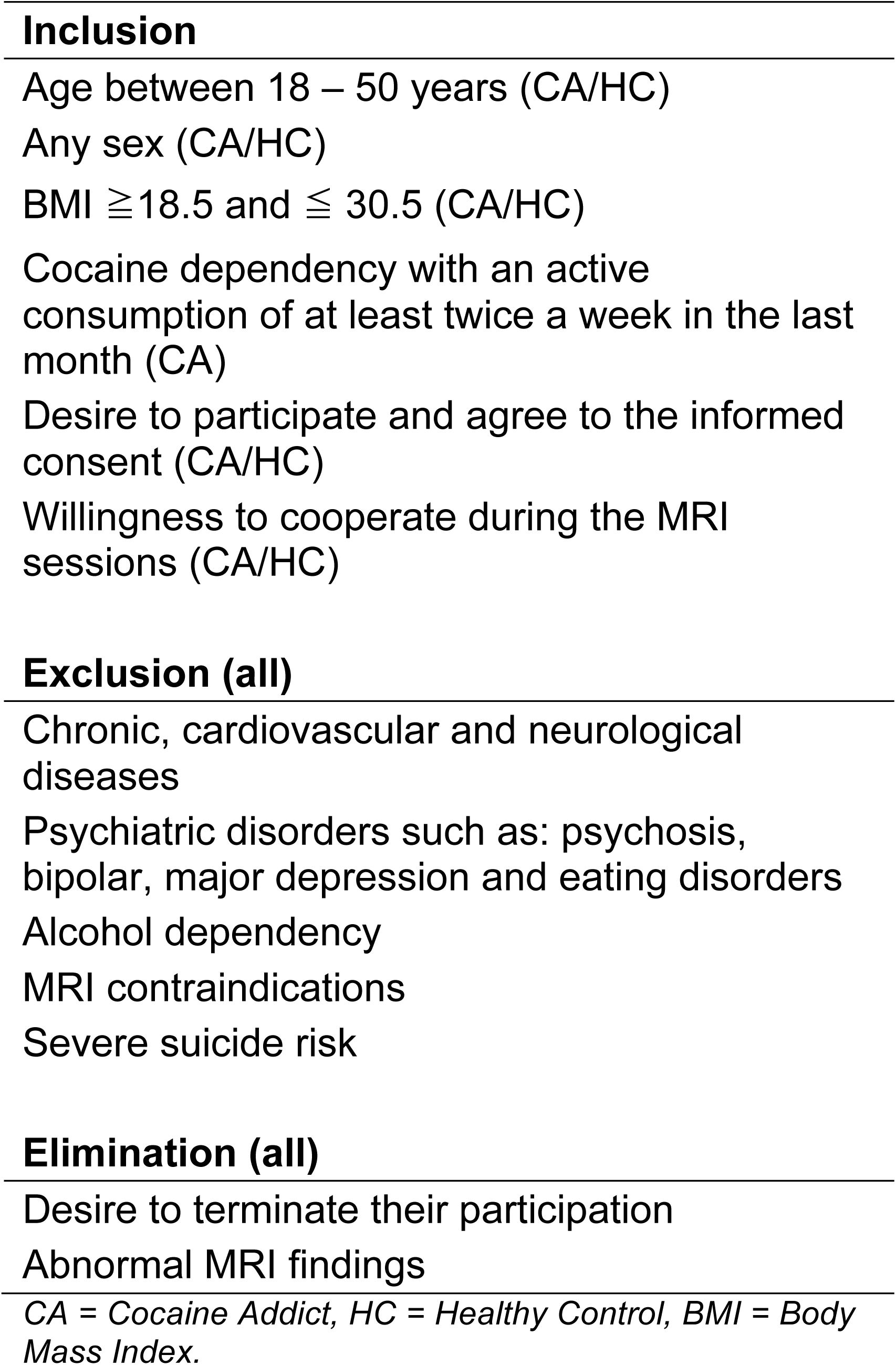
Recruitment criteria.

**Supplementary Table 3.** Clinical Instruments.

| **Instruments** | **For Criteria** |
| --- | --- |
| MINI-Plus | x |
| Addiction Severity Index (ASI) |  |
| Diagnostic Interview for Borderlines-revised, DIB-R |  |
| Clinical Global Impression Scale for Borderline Personality Disorder (CGI-BPD) |  |
| Edinburgh Handedness Inventory |  |
| AMAI NSE 8x7 rule questionnaire |  |
| Structured clinical interview for DSM Axis II, self report (SCID-II) | x |
| Symptom Checklist-90-revised (SCL-90-R) |  |
| Barratt Impulsiveness Scale-11 (BIS-11) |  |
| Difficulties in Emotion Regulation Scale (DERS) |  |
| Timeline follow back |  |
| WHO Disability Assessment Schedule (WHODAS) 2.0 |  |
| Cocaine Craving Questionnaire, general and now (CCQ) |  |
| Demographic data form, self report |  |
| Clinical Interview | x |

For Criteria = Instruments used for inclusion and exclusion criteria.

**Supplementary Table 4.** Psychiatric comorbidities of cocaine addicts.

| **Psychiatric Comorbidities** | **Count (n = 36)** |
| --- | --- |
| Major Depressive Episode current (2 weeks) | 4 |
| Major Depressive Episode Recurrent | 4 |
| Substance Induced Mood Disorder | 7 |
| Major Depressive Episode with Melancholy | 2 |
| Suicidality current (past month) | 13 |
| Suicidality past | 2 |
| Anxiety Disorder with panic due to a general medical condition current | 1 |
| Social Phobia current (past month) | 2 |
| Specific Phobia current | 3 |
| Post-traumatic Stress Disorder current (past month) | 3 |
| Alcoholic dependence (past 12 months) | 13 |
| Alcohol dependence lifetime | 17 |
| Alcohol abuse (past 12 months) | 17 |
| Alcohol abuse lifetime | 22 |
| Substance dependence (non alcohol; past 12 months) | 31 |
| Substance dependence (non alcohol; lifetime) | 32 |
| Generalized anxiety disorder current (past 6 months) | 5 |
| Substance induced generalized anxiety disorder current | 1 |
| Antisocial Personality Disorder lifetime | 10 |
| Somatization Disorder lifetime | 1 |
| Somatization Disorder current | 1 |
| Hypochondriasis current | 1 |
| Conduct Disorder (past 12 months) | 1 |
| Attention Deficit/ Hyperactivity Disorder (past 6 months) | 12 |
| Attention Deficit/ Hyperactivity Disorder (children adolescents; current) | 2 |
| Attention Deficit/ Hyperactivity Disorder (children adolescents; lifetime) | 2 |
| Attention Deficit/ Hyperactivity Disorder (adults; lifetime) | 13 |
| Attention Deficit/ Hyperactivity Disorder (adults; current) | 13 |
| Adjustment Disorders current | 1 |

Comorbidities extracted from the Mini-PLUS interview in Spanish version 5.0.0

**Supplementary Table 5.** Lifetime medication of cocaine addicts.

| **Medication** | **Count (n = 36)** |
| --- | --- |
| Valproic Acid | 6 |
| Fluoxetine | 5 |
| Pregabalin | 3 |
| Carbamazepine | 2 |
| Hydroxyzine | 1 |
| Risperidone | 2 |
| Sertraline | 1 |
| Olanzapine | 1 |
| Alprazolam | 1 |
| Not Specified | 1 |

Based on patients' reports in the general interview and the Addiction Severity Index.

**Supplementary Table 6.**
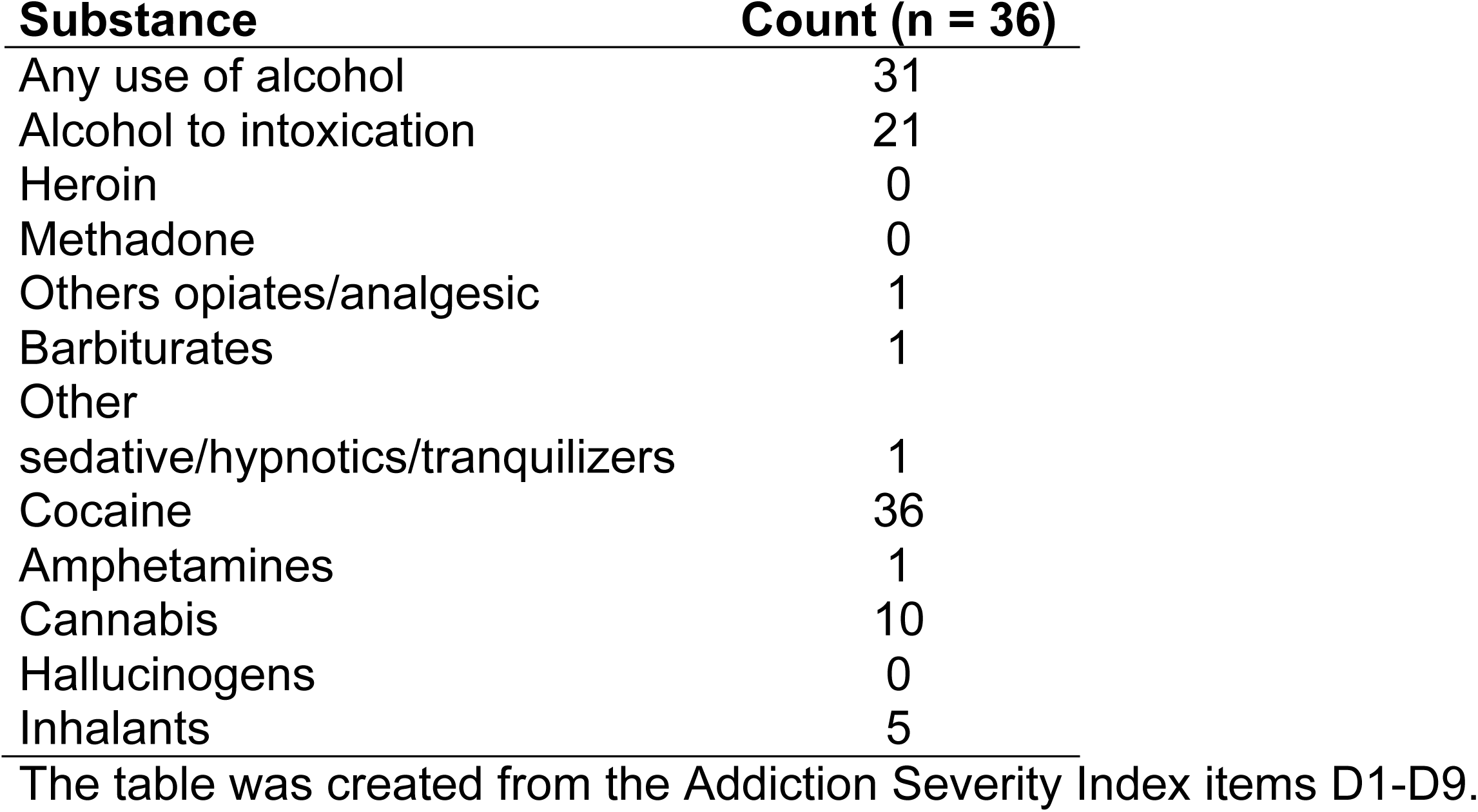
Lifetime history of substance use and abuse.

**Supplementary Table 7.** Regression analysis of Right Pulvinar Volume

|  | **B** | **SE B** | **$\beta$** |
| --- | --- | --- | --- |
| **Step 1** |  |  |  |
| Constant | 2153.74 | 55.75 |  |
| Years of Consumption | -15.96 | 4.59 | -0.52 ** |
| **Step 2** |  |  |  |
| Constant | 771.7 | 406.5 |  |
| Years of Consumption | -14.51 | 4.23 | -0.48 ** |
| Whole Brain Volume | 0.00096 | 0.00033 | 0.44 ** |
| Sex | 51.59 | 194.5 | 0.04 |
| **Step 3** |  |  |  |
| Constant | 1174 | 576.4 |  |
| Years of Consumption | -11.41 | 5.27 | -0.37 * |
| Whole Brain Volume | 0.00082 | 0.00036 | 0.37 * |
| Age | -5.65 | 5.73 | -0.19 |
| Sex | -3.35 | 202.5 | -0.003 |

$R^2 = 0.25$ for Step 1, $\Delta R^2 = 0.43$ for Step 2, $\Delta R^2 = 0.43$ for Step 3, \* = $p < 0.05$ , \*\* = $p < 0.01$ , B = estimate, SE B = Standard Error.

**Supplementary Table 8.**
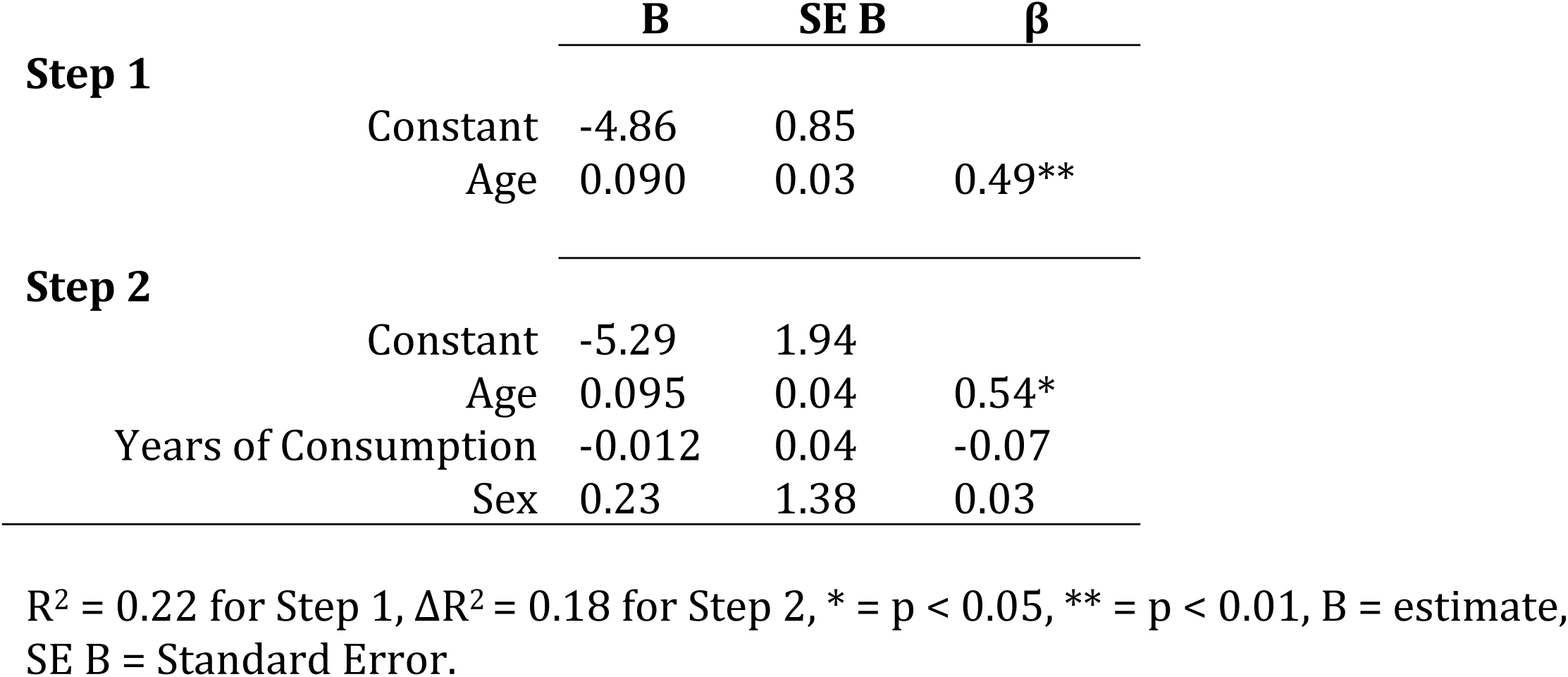
Regression analysis of Left Striatum Displacement Cluster peak

**Supplementary Table 9.** Regression analysis of Left Thalamus MKT.

|  | **B** | **SE B** | **$\beta$** |
| --- | --- | --- | --- |
| **Step 1** |  |  |  |
| Constant | 1.04 | 0.02 |  |
| Years of Consumption | -0.004 | 0.001 | -0.62** |
| **Step 2** |  |  |  |
| Constant | 1.01 | 0.05 |  |
| Years of Consumption | -0.005 | 0.002 | -0.74* |
| Age | 0.001 | 0.002 | 0.18 |

$R^2 = 0.40$ for Step 1, $\Delta R^2 = 0.31$ for Step 2, \* = $p < 0.05$ , \*\* = $p < 0.01$ , B = estimate, SE B = Standard Error.

